# Expanded detection and impact of *BAP1* alterations in cancer

**DOI:** 10.1101/2023.11.21.568094

**Authors:** Ian R. Sturgill, Jesse R. Raab, Katherine A. Hoadley

## Abstract

Aberrant expression of the *BAP1* tumor suppressor gene is a prominent risk factor for several tumor types and is important in tumor evolution and progression. Here we performed integrated multi-omic analyses using data from The Cancer Genome Atlas (TCGA) for 33 cancer types and over 10,000 individuals to identify alterations leading to *BAP1* disruption. We combined existing variant calls and new calls derived from a *de novo* local realignment pipeline across multiple independent variant callers, increasing somatic variant detection by 41% from 182 to 257, including 11 indels ≥40bp. The expanded detection of mutations highlights the power of new tools to uncover longer indels and impactful mutations. We developed an expression-based *BAP1* activity score and identified a transcriptional profile associated with *BAP1* disruption in cancer. *BAP1* has been proposed to play a critical role in controlling tumor plasticity and normal cell fate. Leveraging human and mouse liver datasets, *BAP1* loss in normal cells resulted in lower *BAP1* activity scores and lower scores were associated with a less-differentiated phenotype in embryonic cells. Together, our expanded *BAP1* mutant samples revealed a transcriptional signature in cancer cells, supporting *BAP1’s* influences on cellular plasticity and cell identity maintenance.

## Introduction

*BRCA* associated protein-1 (*BAP1*) has been identified as a tumor suppressor gene with activity across a broad range of biological processes (1–3) including DNA damage repair (1, 4, 5), cell cycle and cell proliferation (1, 6–8), and apoptosis (8–10). *BAP1* mutations promote development of cancer across multiple tissue types and are associated with worse survival outcomes (11–14).

*BAP1* mutations are most prevalent in uveal melanoma, mesothelioma, and renal cell carcinoma (1, 15). Mutations occur very infrequently in other cancer types, often leading to exclusion from consideration due to underpowered analyses. However, among the cancer types that have been studied, consequences of *BAP1* loss are pleiotropic. It is therefore reasonable to assume that there is tissue-specific context that has not yet been systematically explored, presenting an opportunity to expand our understanding of the role of *BAP1* in cancer – particularly if we can incorporate additional cancer types into a larger set of analyses of *BAP1* alteration.

Although *BAP1* mutations have been studied in a small subset of cancer types (1, 11–15), there has been limited incorporation of other types of alterations that can lead to *BAP1* loss. In mesothelioma, we demonstrated an increase of 40% of alterations leading to decreased *BAP1* expression by accounting for copy number loss and previously-undetected long indels (16). This suggests that many prior studies which focused more on single-nucleotide variants have missed important subsets of *BAP1*-altered samples, thereby impacting interpretation of the role of *BAP1*. Here, we account for these additional types of alterations across a broader set of cancer types represented in The Cancer Genome Atlas (TCGA) to better characterize functional consequences of *BAP1* alteration in cancer. We previously demonstrated that *BAP1* loss characterized a cholangiocarcinoma-like (CHOL-like) expression-based subset of liver hepatocellular carcinomas and hypothesized that CHOL-like liver tumors represented *BAP1*-driven changes resembling a transdifferentiated cell phenotype (14). To understand the specific impact of *BAP1* loss both in cancer and normal biology, we developed an RNA expression-based *BAP1* activity score and validated this score in external human and mouse liver datasets. We linked this activity score to embryonic development in murine liver supporting a role for *BAP1* driving changes in phenotypic plasticity and cell states.

## Materials and Methods

### Somatic mutation calling workflow with *de novo* local realignment

10,414 pairs of tumor and matched-normal aligned binary alignment map (BAM) slice files of the *BAP1* locus and 100kb flanking regions from 33 cancer types were downloaded from The Cancer Genome Atlas (TCGA) Genomic Data Commons (GDC)’s harmonized hg38 mapped data using the command line GDC Data Transfer Tool v.1.6.1. The specific genomic region was hg38 chr3:52300003-52511030. Individual BAM slices were then processed through a Nextflow DSL 2 pipeline for somatic mutation calling. Briefly, slices were indexed using Samtools (version 1.9) (17). Reads were realigned to the hg38 human reference genome (GRCh38.d1.vd1) using the ABRA2 (version 2.24-0) *de novo* local realigner with --undup and --no-edge-ci parameter flags (18). Somatic mutations were called using the Strelka2 somatic workflow (version 2.9.10) and Cadabra indel (version 2.24-0) variant callers after marking duplicate reads using bammarkduplicates from biobambam (version 2.0.87) (18–20). Raw variant call format (VCF) output files were converted to mutation annotation format (MAF) using vcf2maf (version 1.6.21) and VEP (cache version 103) (21, 22).

### Filtering, manual review, and characterization of somatic variants

Variant calls were initially filtered based on individual variant caller quality tags (i.e., FILTER == PASS). These variant calls were merged with calls from the GDC’s internal mutation calling pipeline consisting of four variant callers: MuSE, MuTect2, VarScan2, and pindel. Variants were further filtered to those that had variant allele frequency (VAF) ≥0.2 and mutant allele read count (t_alt_count) ≥2 followed by manual review using the Integrative Genomics Viewer (IGV) to further assess read support strength and evidence of any other confounding factors such as alignment at low-complexity regions and variant bases captured only by ends of reads (23). Current variant calls for *BAP1* were then compared to historical calls generated for TCGA in the MC3 dataset and characterized by their somatic MAF variant classification (24). Chromosomal start locations of mutations were plotted, together with variant classification information, on a lollipop plot adapted from cBioPortal’s MutationMapper web tool (25, 26). Protein-level domain annotations and visualization were approximated and adapted from Haugh et al. (27).

### Definition of copy number loss

Estimates of focal gene-level copy number for *BAP1* and other genes were downloaded from the GDC, which used the hg38 ABSOLUTE LiftOver workflow to provide per-gene integer value estimations of gene copy numbers for each sample (28, 29). Samples with gene-level copy number of less than two corresponded to copy number loss. Segment widths of samples with copy number loss were computed from segment start and end loci from masked copy number segment files from the GDC, which provided estimates of genomic windows of segments containing the *BAP1* gene locus using the DNAcopy workflow (30). Plots of segments in relation to the chromosome were generated using plotgardener (version 1.10.2) (31).

### Differential mRNA expression and gene set enrichment analyses (GSEA)

Per-sample hg38 Spliced Transcripts Alignment to a Reference (STAR)-aligned unstranded counts files were downloaded from the GDC (32). Where appropriate, we adjusted for additional sources of variation (tumor purity, histological subtype, and/or previously-defined molecular subtype) in our model design in a cancer type-specific manner (Supplemental Table S3B). Samples without sufficient subtype annotations and samples without tumor RNA sequencing data available were excluded from analyses. Sample groups were then tested for differential gene expression using the DESeq2 R package (version 1.44.0) (33). Briefly, we performed Wald testing with independent hypothesis weighting provided by the IHW R package (version 1.32.0) (34).

Initial results were then subjected to adaptive shrinkage estimation using the apeglm R package (version 1.26.1) (35). Genes were annotated using biomaRt (version 2.60.1) via the annotables R package (version 0.2.0) (36–38). Gene set and pathway analyses were conducted using the fgsea R package (version 1.30.0) and data from the C8 gene sets from the MSigDB database (39, 40). Volcano plots were generated using the EnhancedVolcano R package (version 1.22.0) and heatmaps were generated using the ComplexHeatmap R package (version 2.20.0) (41, 42).

### Gene set and *BAP1* activity scores

*BAP1* activity scores were computed based on a gene set of differentially expressed genes between altered-with-mutation (Mut and Mut+CN) and unaltered tumors for five cancer types: cholangiocarcinoma (CHOL), kidney renal cell clear cell carcinoma (KIRC), liver hepatocellular carcinoma (LIHC), mesothelioma (MESO), and uveal melanoma (UVM). Expression values of genes upregulated in *BAP1*-mutant tumors were summed and multiplied by -1, while expression values of downregulated genes were summed and multiplied by +1. These per-sample values were then summed. We used the mclust (version 6.1.1) R package with default parameters to perform model-based clustering of activity scores and classify samples as mutant-like or wildtype-like (43). This scoring was then applied to all samples in TCGA. For the bile duct gene signature, gene sets BILE_DUCT_CELLS_1 through 4 from the MSigDB C8 cell type signature database were combined (40). Bile duct signature scores were computed as z-scores of median-centered, log2-transformed counts of genes from the combined gene set.

### Statistical Analysis

Statistical analysis was performed using functions included in R version 4.4.1. Specific methods are listed alongside their corresponding results.

## Results

### *BAP1* somatic mutations in TCGA are underestimated and are predominantly deleterious

To determine the extent of underestimation of *BAP1* mutations in TCGA and add occurrences of larger indels to existing mutation calls, we aligned BAM slice files containing the *BAP1* locus and 100kb flanking regions to the hg38 genome using a *de novo* local realignment approach with ABRA2 and two variant callers: Cadabra and Strelka2. These variant calls were merged with updated TCGA GDC calls which were also aligned to the hg38 genome and generated from four independent variant callers: MuSE, MuTect2, VarScan2, and Pindel. Out of 10,414 samples and 33 cancer types, a total of 1337 non-synonymous, non-intronic variants (1329 somatic and 8 germline) were detected in 988 individuals across the two updated hg38-aligned approaches, including 14 indels ≥40bp in length (Supplemental Table S2A). We conducted further review of variants to select those with greater support (tumor variant allele frequency ≥0.2, tumor alternate allele read count ≥2, and manual review of read alignment in IGV), resulting in a reduction from 1337 to 265 variants in 233 individuals (2.2%) across TCGA cancer types, including 11 larger indels ≥40bp that were not previously detected (Figure 1A-B and Supplemental Table S2C).

**Figure 1.**
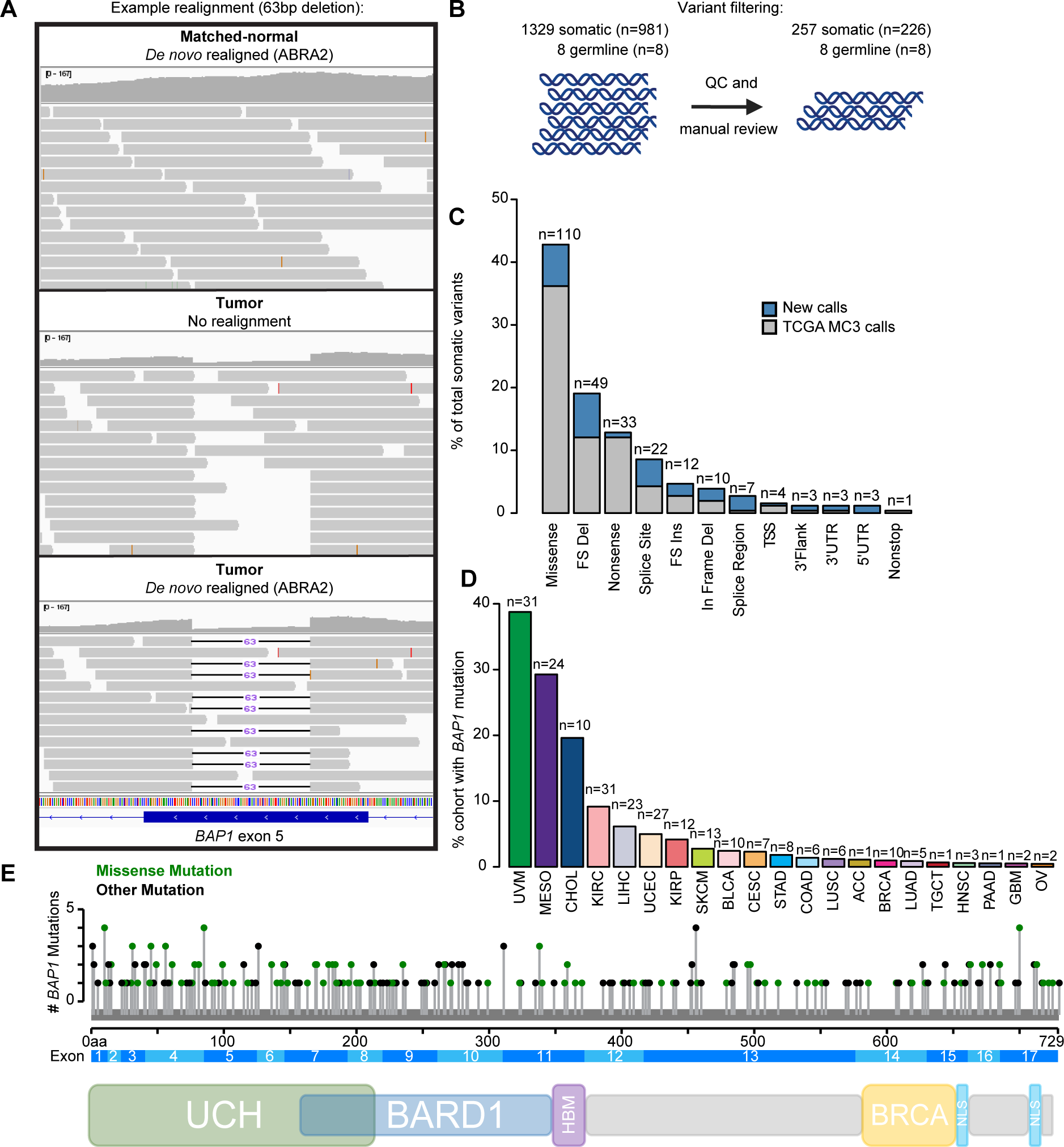
Prevalence and characterization of *BAP1* mutations in TCGA pan-cancer tumor types. **(A)** Example *de novo* realignment with ABRA2 to the hg38 reference genome for a 63-bp indel detected in an LIHC tumor sample (TCGA-FV-A3I0) with visualization in IGV. **(B)** Schematic indicating numbers of *BAP1* variants before and after filtering and manual review. Created with BioRender.com. **(C)** Distribution of *BAP1* somatic variant types (total=257 variants). Blue, additional new mutations detected in the present workflow combined with updated TCGA GDC variant calls; grey, prior TCGA MC3 mutation calls. **(D)** Frequency of *BAP1* mutant tumors by TCGA tumor type. **(E)** Lollipop plot of *BAP1* somatic variant locations across the length of the gene, adapted from cBioPortal (25, 26). Green, missense mutations; black, all other mutation types. Bottom: protein-level schematic of the major domains of interest, adapted from Haugh et al. (27). UCH: ubiquitin carboxy-terminal hydrolase, BARD1: *BARD1* binding domain, HBM: *HCF* binding motif, BRCA1: *BRCA1* binding domain, NLS: nuclear localization signal.

We compared somatic variants (257/265 total variants) to an earlier iteration of variant calling used to generate the TCGA MC3 dataset, which included alignment to the hg19 genome and seven callers: MuTect, Varscan2, Indelocator, Pindel, SomaticSniper, RADIA, and MuSE (24). This earlier pipeline resulted in the detection of 251 variants from 224 individuals, of which 190 variants from 170 individuals similarly passed simple call quality filtering, none of which were ≥40bp (Supplemental Figure S1 and Supplemental Table S2B and S2D). Of the 182 somatic variants in MC3, 77 (30%) were new calls and the remaining 180 were concordant (Figure 1C). All MC3 variants and mutant samples were captured in the new approach, except one variant, a 5bp deletion in KIRP sample TCGA-2Z-A9JD (Supplemental Tables S2C and S2D). However, our realignment approach detected two single-nucleotide variants (deletion and missense) in the similar region, suggesting a potential difference in hg19 versus hg38 alignment and downstream variant calling.

The majority of *BAP1* somatic variants were missense mutations (110/257, 42.8%), followed by deleterious nonsense and frameshift combined events (94/257, 36.6%, Figure 1C and Supplemental Table S2C). Of the cancer types represented in TCGA, 21/33 (63.6%) have at least one *BAP1* mutant sample (Figure 1D). Six tumor types had >5% mutation frequency: cholangiocarcinoma (CHOL), kidney clear cell renal cell carcinoma (KIRC), liver hepatocellular carcinoma (LIHC), malignant mesothelioma (MESO), uterine corpus endometrial carcinoma (UCEC), and uveal melanoma (UVM) (Supplemental Table S3A).

Somatic variants were broadly distributed across the length of the gene with no significant difference in protein-level domain representation for missense versus other mutation types (two-sided Fisher’s exact test p=0.1) (Figure 1E and Supplemental Table S2C). We observed 161/257 (62.6%) variants occurred at an annotated functional domain with 110/257 (42.8%) occurring at the catalytic UCH domain (Figure 1E and Supplemental Table S2C). Due to relatively small numbers of variants, we were insufficiently powered to further explore potential differences in impacts to specific functional domains.

### Copy number alterations constitute a major mechanism of *BAP1* loss across cancer types

To assess the contribution of copy number (CN) alterations to a *BAP1* loss phenotype, we examined gene-level CN estimates of the *BAP1* gene. *BAP1* CN loss was observed in 1547 samples and estimated at single copy loss in 99% of those samples (Table S3A-B). Gene-level CN loss occurs frequently (>30% of samples) in KIRC, UVM, CHOL, MESO, lung squamous cell carcinoma (LUSC), and head and neck squamous cancers (HNSC) (Figure 2A and Table S3A).

**Figure 2.**
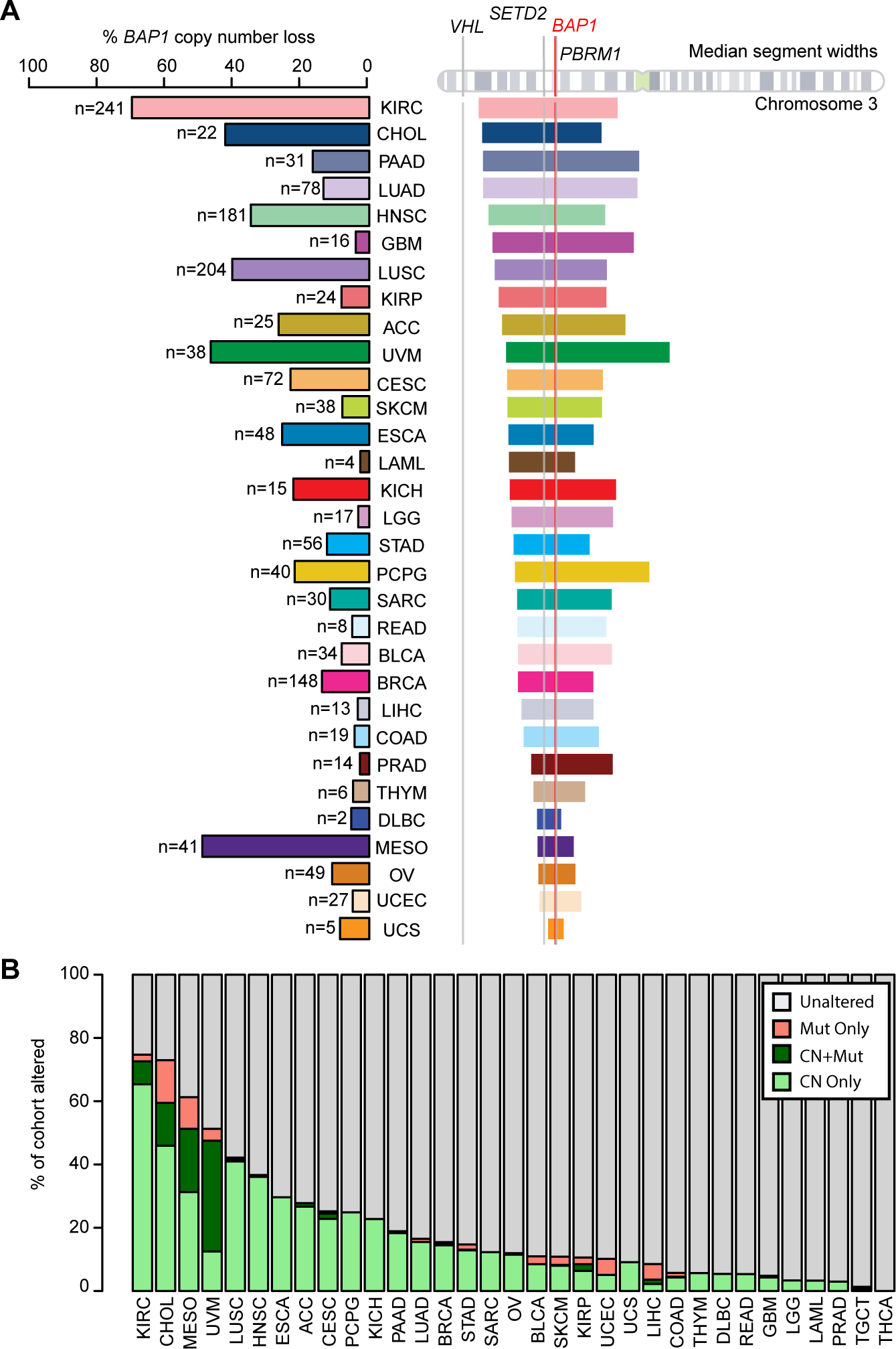
Copy number loss of *BAP1* is frequent in cancer. **(A)** Left: percent and number of samples with *BAP1* gene-level copy number loss for each cancer tumor type. Right: representations of median segment widths of CN loss in samples with *BAP1* gene-level CN loss aligned against chromosome 3, which contains the indicated *BAP1* gene locus. The locations of other commonly altered driver genes in cancer (*VHL, SETD2,* and *PBRM1*) are also indicated. **(B)** Percentage of each tumor type altered for *BAP1*, colored by specific alteration type. Mut Only: mutation only, CN+Mut: co-occurring gene-level copy number loss and mutation, CN Only: gene-level copy number loss only.

We also considered the CN segment widths around *BAP1* to determine if the loss is focal or arm-level. For many cancer types, the loss is arm-level, spanning large regions of the ∼90Mb chromosome 3p arm on which *BAP1* is located and including regions corresponding to over 1000 annotated genes. There are other commonly lost tumor suppressors on this chromosome (*VHL*, *SETD2*, and *PBRM1*). We looked at the width of the deletion segment by tumor type and found variation by tumor type. For example, KIRC (median 60.7Mb, interquartile range IQR 33.6-82.3Mb), LUAD (median 63.0Mb, IQR 34.6-87.8Mb), and PCPG (median 40.8Mb, IQR 12.3-89.2Mb) have very large segment widths of loss (Supplemental Table S3B). In contrast, MESO (median 8.3Mb, IQR 0.2-29.2Mb), OV (median 9.5Mb, IQR 4.7-19.4Mb), and UCEC (median 7.2Mb, IQR 4.6-19.4Mb) have relatively more focal *BAP1* CN loss, impacting an estimated 200-360 genes.

Combining CN and mutation data, 1679 samples from 32 tumor types were considered altered (Figure 2B and Supplemental Table S3A-S3B). Alterations were predominantly copy number-driven (1547/1679, 92.1%). Mutations were less frequent; they occurred alone in 7.9% (132/1679) of altered samples and co-occurred with estimated single-copy CN loss in 6% (101/1679). Regardless of tumor purity, mean variant allele frequency is 0.46 ± 0.21 (standard deviation). Variant allele frequency is higher in samples with both mutations and copy number loss compared to mutations alone (two-sided Mann-Whitney U test with continuity correction p=0.002, Figure S2A). Variant allele frequency and tumor purity have a moderate positive correlation (linear regression Wherry adjusted r^2^=0.35, Figure S2B), suggesting that these mutations are less likely to be subclonal. The most highly-altered cancer types for *BAP1* are KIRC, CHOL, UVM, MESO, LUSC and HNSC – all with >30% altered samples within their tumor types (Figure 2B and Supplemental Table S3A-S3B). Among altered samples, there is also substantial variability in the mechanism of alteration. In KIRC and HNSC, alterations are primarily driven by instances of CN loss. However, in CHOL, UVM, MESO, and LIHC, mutations appear to play a much larger role. All alteration types, regardless of cancer type, resulted in lower RNA-level expression of *BAP1* than unaltered samples (two-sided pairwise Mann-Whitney U test with continuity correction p<0.01 for each alteration type, Figure S3 and Supplemental Table S3B).

### Expression-based signatures identify a distinct *BAP1* mutant-like subset

To better understand the tissue and context-specific consequences of *BAP1* alteration, we examined differential gene expression across samples with a *BAP1* mutation (*BAP1* mutation and *BAP1* mutation plus copy number loss) and those without any alteration (no mutation or copy number loss) by tumor type among five tumor types with higher >5 *BAP1* mutant samples: CHOL, KIRC, LIHC, MESO, and UVM. We reasoned that comparing *BAP1* mutant to unaltered samples would give the most accurate *BAP1*-dependent gene expression changes. We adjusted for additional sources of known variation such as tumor purity and subtype in our model designs in a cancer-specific manner (Supplemental Table S3B). Of 16,333 expressed protein coding genes tested in this manner, 3966 (24.3%) were significantly changed at an adjusted p-value less than 0.05 (Figure 3A and Supplemental Table S4A-S4B).

**Figure 3.**
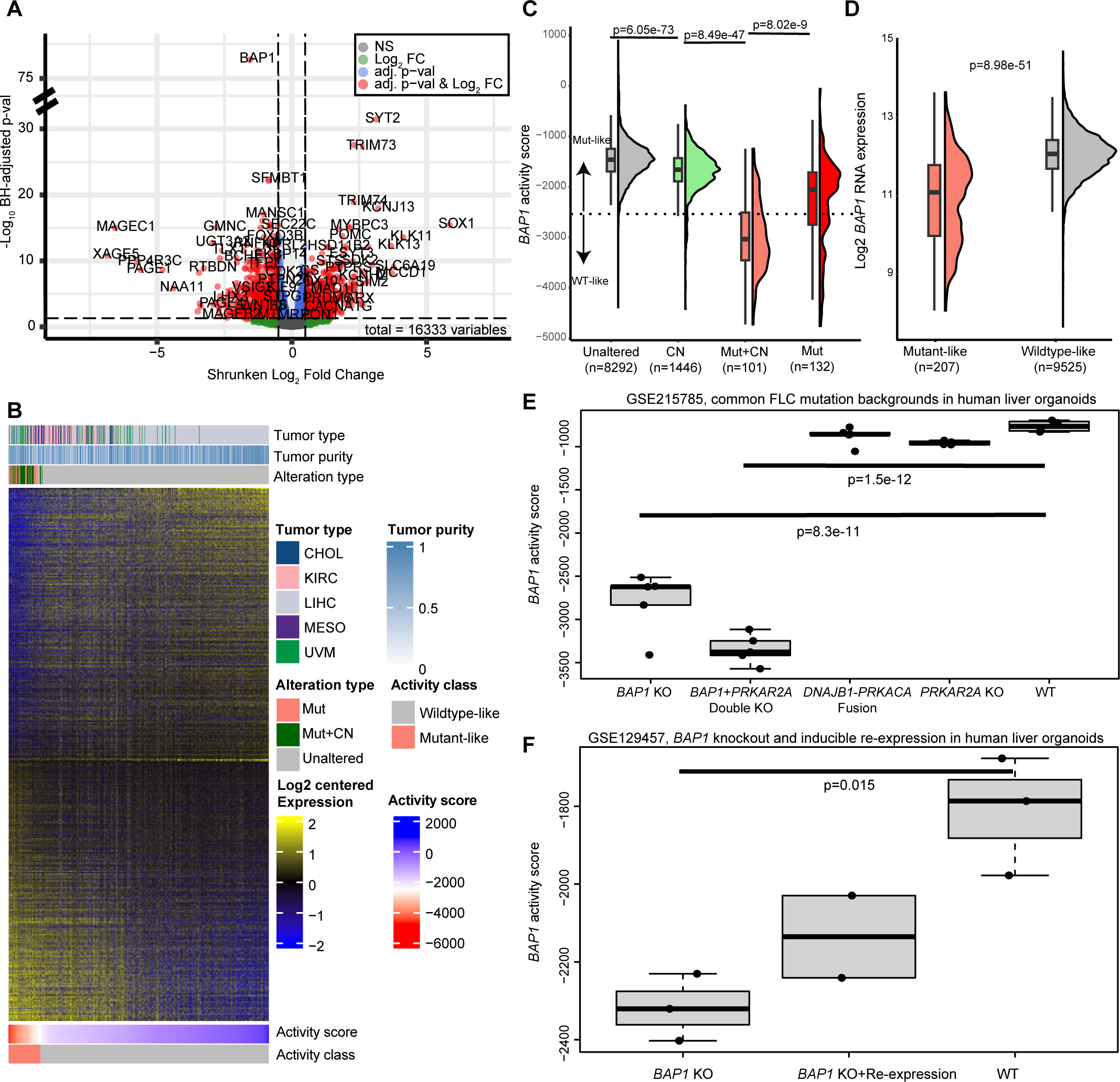
*BAP1* activity scores capture *BAP1*-driven changes in gene expression across multiple independent datasets. **(A)** Volcano plot of *BAP1* differentially expressed genes between *BAP1-*mutant and unaltered tumors from TCGA CHOL, KIRC, LIHC, MESO, and UVM tumor types. Adjusted p-values are derived in DESeq2 using the Benjamini-Hochberg (BH) procedure. Log_2_ fold change values were shrunken using the apeglm R package. Red: adjusted p-value<0.05 and absolute log_2_ fold change≥1.5, blue: adjusted p-value<0.05, green: absolute log_2_ fold change≥1.5, grey: not significant (NS). **(B)** Heatmap of differentially expressed genes between *BAP1*-mutant and unaltered tumors. Mut: mutation only, Mut+CN: mutation and gene-level copy number loss. **(C)** TCGA pan-cancer activity scores across the *BAP1* alteration types. The dotted line represents the threshold delineating wildtype-like and mutant-like classification. CN: copy number loss only, Mut+CN: mutation and gene-level copy number loss, Mut: mutation only. P-values are derived from pairwise two-sided Mann-Whitney U testing with continuity correction and Bonferroni adjustment. **(D)** TCGA pan-cancer comparison of *BAP1* RNA expression in wildtype-like and mutant-like tumors. P-value derived from two-sided Mann-Whitney U testing with continuity correction and Bonferroni adjustment. **(E)** Comparison of *BAP1* activity scores for a dataset of human liver organoids (GSE215785) with experimentally-produced common fibrolamellar carcinoma mutation backgrounds. *BAP1* KO: *BAP1* single-knockout, *BAP1*+*PRKAR2A*: *BAP1* and *PRKAR2A* double-knockout, *PRKAR2A* KO: *PRKAR2A* single-knockout, WT: wildtype. P-values are derived from pairwise two-sided t-test with Bonferroni adjustment. **(F)** Comparison of *BAP1* activity scores for a dataset of human liver organoids (GSE129457) with conditional *BAP1* knockout. *BAP1* KO: *BAP1*-knockout, *BAP1* KO+Re-expression: *BAP1* knockout with 24-hour re-expression, WT: wildtype. P-value derived from pairwise two-sided t-test with Bonferroni adjustment.

Using this set of differentially expressed genes, we computed per-sample *BAP1* mutation signature scores and classified tumors as being more mutant-like or wildtype-like (see Methods). We ordered tumors by their respective scores, revealing a distinct pattern of gene expression among mutant-like tumors (Figure 3B). Among the five tumor types in this analysis, 57/59 (96.6%) of mutant tumors were classified as mutant-like in this manner. We then applied this mutation signature scoring method to the rest of the TCGA dataset, where 117/233 (50.2%) of mutant tumors were classified as mutant-like, with mutant tumors having lower activity scores and tumors with *BAP1* mutation plus copy number loss having the lowest scores (Figure 3C). We further confirmed that mutant-like tumors had lower RNA-level expression of *BAP1*, consistent with a signature driven by *BAP1* loss (Figures 3D).

To further verify that our *BAP1* activity signature effectively captures a transcriptional profile shift associated with *BAP1* loss, we leveraged two external datasets of *BAP1* knockout in human liver organoids. In the first dataset (GSE215785), common fibrolamellar carcinoma mutant backgrounds were generated, including *BAP1* single-knockout and *BAP1*/*PRKAR2A* double-knockout organoids (44). Both single- and double-knockout organoids had significantly lower *BAP1* activity scores relative to wildtype (pairwise t-test with Bonferroni correction p=8.3e-11 and p=1.5e-12 respectively, Figure 3E), but no difference was observed in the samples with the classic *DNAJB1-PRKACA* fusion or *PRKACA* knockout which is the known major driver of that subset of liver cancer. In the second dataset (GSE129457), *BAP1* knockout organoid models were generated both with and without *BAP1* re-expression (45). Consistent with GSE215785 data in Figure 3E, we also observed reduced *BAP1* activity scores among *BAP1* knockout organoids relative to wildtype (pairwise t-test p=0.015, Figure 3F). Re-expression of *BAP1* in the knockout organoid led to increased activity scores relative to knockout (Figure 3F).

### *BAP1* alteration characterizes a molecularly distinct subset of hepatocellular carcinomas

Because *BAP1* alterations have been previously shown to constitute a molecularly distinct subset of tumors in LIHC and may play a unique role in regulating cell identity and differentiation processes in the liver, we selected LIHC for more focused analyses (14, 44–46). In LIHC, *BAP1* mutant-like tumors were distinct from wildtype-like tumors (Figure 4A). Annotations of tumor molecular subtyping from Damrauer et al. revealed that the *BAP1-*altered subset was enriched for blast-like and cholangiocarcinoma-like (CHOL-like) LIHC samples (14). Consistent with these previous data, 17/31 (54.8%) of *BAP1*-altered tumors were one of these two molecular subtypes; among the unaltered tumors, 77/327 (23.5%) were one of these subtypes. Blast-like and CHOL-like tumors have *BAP1* activity scores lower than the other LIHC tumors (pairwise two-sided Mann-Whitney U test with continuity correction and Bonferroni adjusted p=2.47e-3 and p=2.38e-9 respectively, Figures 4A-B and Supplemental Table S3B).

**Figure 4.**
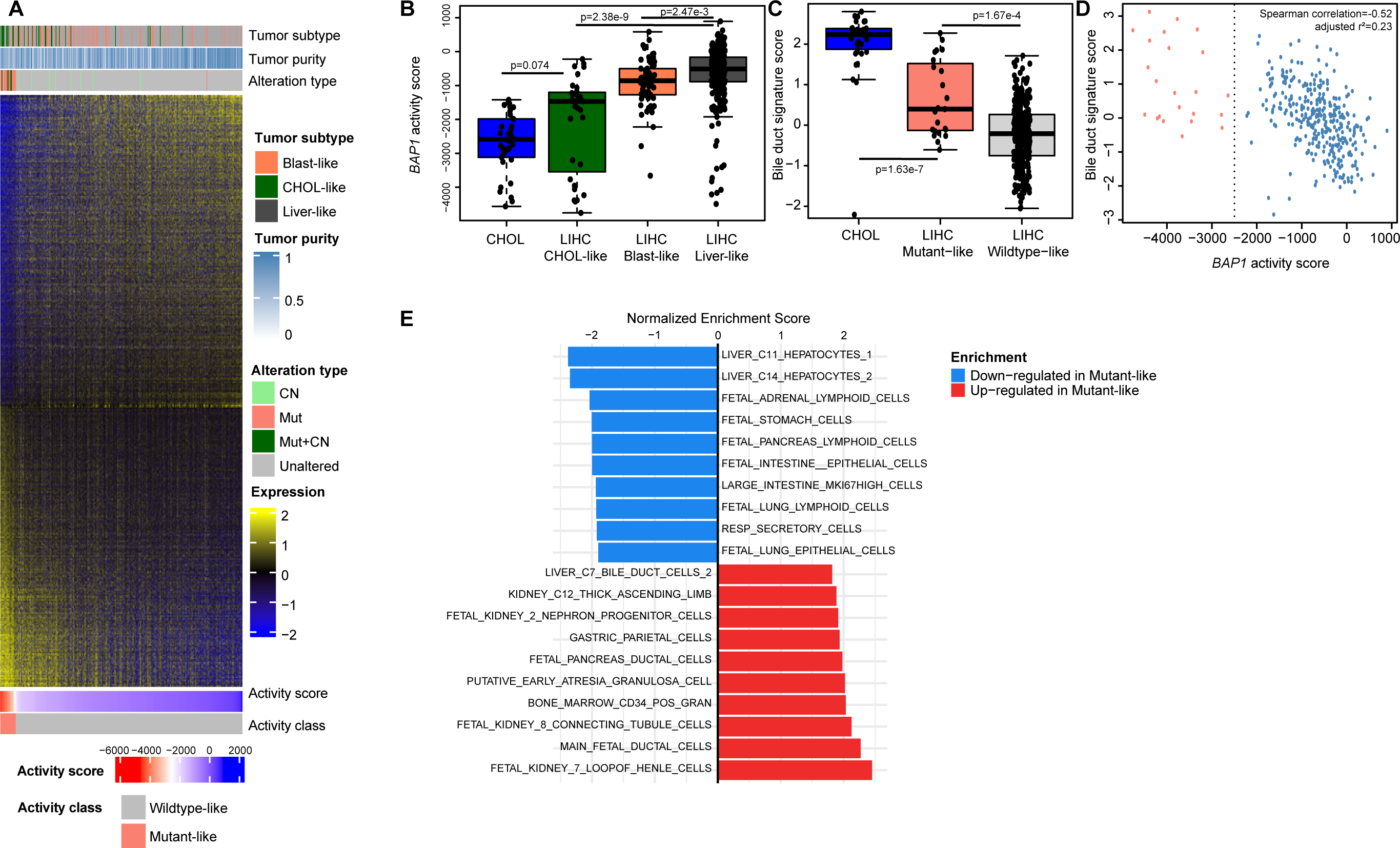
*BAP1* mutant-like hepatocellular carcinoma (LIHC) tumors are associated with a more duct-like phenotype. **(A)** Heatmap of differentially expressed genes between *BAP1* mutant-like and wildtype-like tumors in LIHC with sample annotations. CN: gene-level copy number loss, Mut: mutation only, Mut+CN: mutation and gene-level copy number loss. **(B)** CHOL and LIHC *BAP1* activity scores separated by molecular subtype from Damrauer et al.(14) P-values are derived from pairwise two-sided Mann-Whitney U tests with continuity correction and Bonferroni adjustment. **(C)** Specific comparison between CHOL and LIHC *BAP1* mutant-like and wildtype-like samples of bile duct gene expression from four combined signatures from the MSigDB C8 gene set collection. Adjusted p-values are derived from pairwise two-sided Mann-Whitney U tests with continuity correction and Bonferroni adjustment. **(D)** Scatterplot of *BAP1* activity and bile duct signature scores with indicated Spearman correlation; linear regression of *BAP1* activity and bile duct signature scores with Wherry adjusted r^2^ value also shown. Red points indicate *BAP1* mutant-like samples and blue points indicate *BAP1* wildtype-like samples. Dotted line represents identity (y=x) line. **(E)** Top ten differential cell type signatures from the MSigDB C8 gene set collection in both directions between *BAP1* mutant-like and wildtype-like LIHC tumors. Shown results are statistically significant with Benjamini-Hochberg adjustment. Red: up-regulated in *BAP1* mutant-like samples, blue: downregulated in *BAP1* mutant-like samples.

We previously hypothesized that the CHOL-like subset arose due to de-differentiation (14). To further explore the relationship between *BAP1* and less-differentiated (or more CHOL-like) LIHC tumors, we examined expression of bile duct cell markers based on four combined bile duct cell signature gene sets. We used true CHOL samples as a comparison and observed that *BAP1* mutant-like LIHC tumors were enriched for bile duct markers relative to wildtype-like samples (pairwise two-sided Mann-Whitney U test with continuity correction and Bonferroni adjusted p=1.67e-4, Figure 4C). Further, bile duct signature scores in LIHC tumors had a moderate negative correlation to *BAP1* activity scores (Spearman correlation=-0.52, Figure 4D), suggesting that the signature captures expression of some genes that separate cholangiocyte and hepatocyte identity. A broader analysis of cell type signatures showed enrichment of bile duct and progenitor markers and a decrease in terminally-differentiated cell markers – for example, hepatocyte markers (Figure 4E) -- consistent with a transcriptional profile shift towards a less- or trans-differentiated phenotype.

### Association of *BAP1* with developmental timing and differential chromatin accessibility

To further understand the potential role of *BAP1* in liver development and cellular differentiation, we explored temporal changes in *BAP1* activity scores within an external single-cell RNA-seq dataset of mouse embryonic liver development (GSE90047) (47). Principal component analysis of single-cell data demonstrates separation of cells both by embryonic timepoint and by putative cell type along the first two principal components (Figure 5A). When *BAP1* activity score values were overlayed on top of these data, there was a clear trend of lower scores (more mutant-like) in cells that are earlier in development; as the embryonic timepoint increases towards a more differentiated phenotype, scores increase along a pronounced gradient (Figure 5B). This trend holds in bulk RNA-seq data from the same study, showing a high correlation between embryonic timepoint and *BAP1* activity score (adjusted r^2^=0.81, Figure 5C).

**Figure 5.**
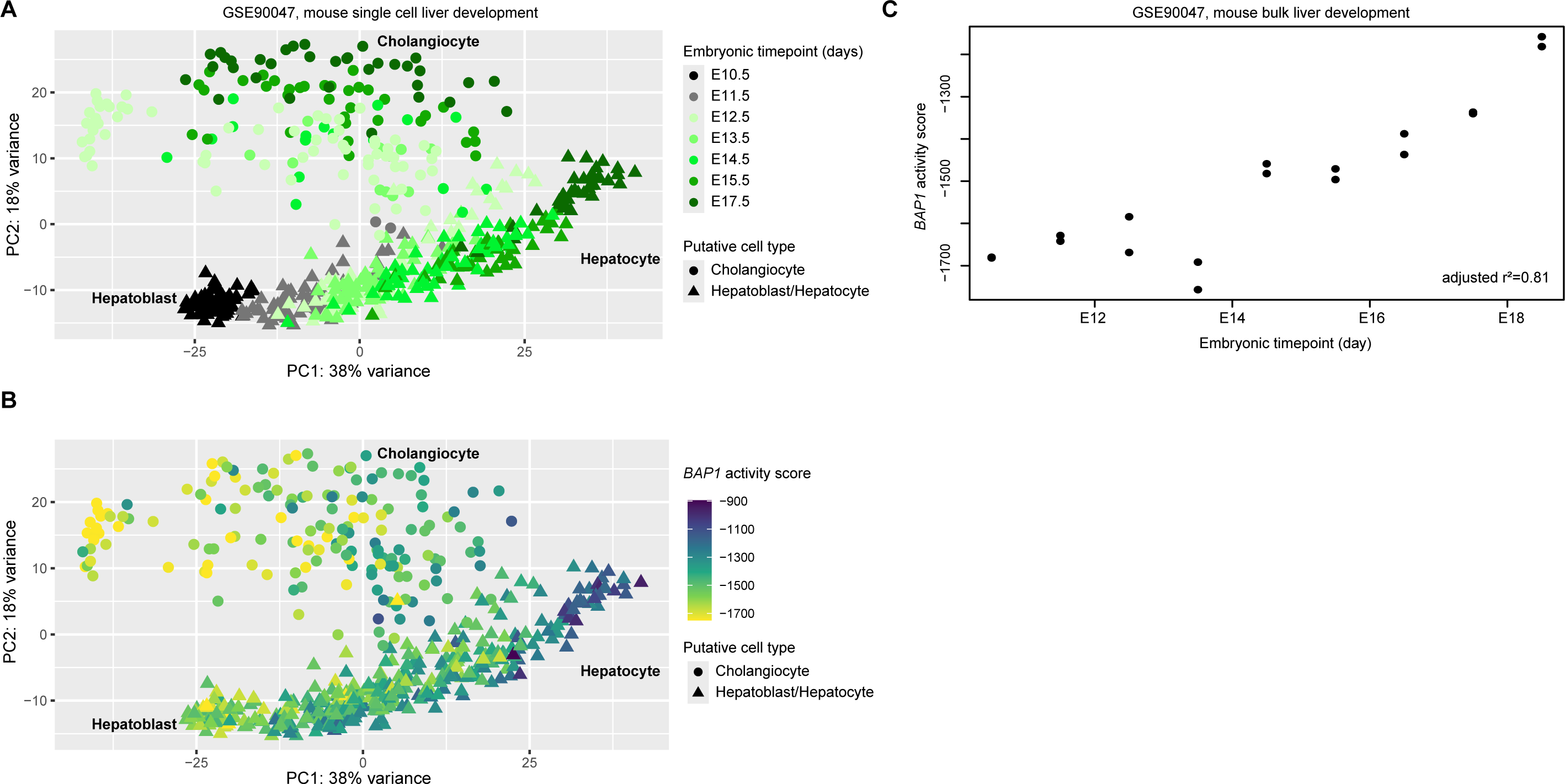
Earlier timepoints in mouse embryonic liver development are associated with lower *BAP1* activity scores. **(A)** Principal component analysis of single-cell RNA-seq data demonstrating temporal separation of putative hepatoblast, hepatocyte, and cholangiocyte cell types, colored by embryonic timepoint. **(B)** Principal component analysis of the same single-cell dataset as A, colored by *BAP1* activity score. **(C)** Scatterplot of embryonic timepoint and *BAP1* activity score in a bulk RNA-seq dataset for mouse liver embryogenesis. Statistic is linear regression with Wherry adjusted r^2^.

## Discussion

### *BAP1* alterations are frequent in cancer

Here we report a 30% increase in detection of *BAP1* somatic mutations across TCGA compared to prior mutation calls (24) by using a *de novo* local realigner and merging variant calls from six individual callers across two pipelines. Importantly, we were able to identify additional indels ≥40bp that previously were undetected. While long read sequencing technology has become more accessible and has improved detection of larger indels, historical datasets like TCGA with short read sequencing will continue to benefit from reanalysis using newer and improved tools capable of detecting additional variants.

Copy number (CN) loss of *BAP1* is found in approximately 20% of TCGA pan-cancer samples, making it the most common alteration of *BAP1* in this cohort (16). However, chromosome 3 is home to multiple tumor suppressors important in many cancers (*VHL*, *SETD2*, *PBRM1*) in addition to *BAP1,* making it unclear in which cancers *BAP1* loss is biologically important. Using gene expression data from tumors with the highest frequency of *BAP1* mutations, we developed a score that could infer which alterations were likely affecting *BAP1* activity or function. We validated our score using data from *BAP1* knockout cancer organoid models which had lower *BAP1* activity scores compared to parental organoids. When we assessed tumors using this score, we found a subset of CN loss events that were associated with reduced *BAP1* activity. Further, *BAP1* CN loss in conjunction with *BAP1* mutations showed the strongest reduction in inferred *BAP1* activity or function.

### Role of *BAP1* in cell identity and differentiation

*BAP1* is important in cell differentiation and development (44, 48–50). Patel and Yanai have postulated that tumor plasticity is “constrained by the organism’s developmental map” (51). For example, hepatocytes and cholangiocytes may both de-differentiate toward their shared progenitor phenotype in tumorigenesis. Our prior work identified as subset of pathologically defined hepatocellular carcinoma (CHOL-Like) with mutations and gene expression profiles more similar to cholangiocarcinoma (14). These tumors were originally defined by *IDH1* loss and we further showed them to have *BAP1* loss as a major altered feature. In our current study, Hepatocellular carcinoma tumors with low *BAP1* activity scores had higher expression of bile duct signatures. Additionally, the subset of CHOL-like hepatocellular carcinoma had lower *BAP1* activity scores more similar to cholangiocarcinoma than to other hepatocellular carcinoma samples.

Our data in tumors suggests that loss of *BAP1* leads to de-differentiation and regulates cell fate. *BAP1* has well-known roles in normal development (48). We assessed changes of *BAP1* activity scores during normal mouse liver development. Using a single-cell RNA-seq mouse embryonic time course, we found that *BAP1* activity scores were low in progenitor cells and increased during liver development (47). These results are consistent with *BAP1’s* role as an important coordinator in the differentiation processes and maintenance of cell identity and support the hypothesis that *BAP1* dysregulation in somatic cells can lead to de-dedifferentiation and cancer.

## Data Availability

### Whole exome sequencing BAM slice files

Because whole exome sequencing BAM files may contain personally identifiable information, they are subject to controlled access through the NIH database of Genotypes and Phenotypes (dbGaP, https://www.ncbi.nlm.nih.gov/gap/). Supplemental Table S1 contains a list of case and file IDs downloaded from the NIH Genome Data Commons repository (https://portal.gdc.cancer.gov/repository).

### Prior MC3 variant calls, tumor purity, and curated survival data

The following data were downloaded from the TCGA PanCanAtlas Publications webpage (https://gdc.cancer.gov/about-data/publications/pancanatlas):

- Pan-cancer variants MAF: mc3.v0.2.8.PUBLIC.maf.gz
- ABSOLUTE purity/ploidy file: TCGA_mastercalls.abs_tables_JSedit.fixed.txt

### Tumor subtyping

Tumor subtype annotations of TCGA samples were compiled from many published sources. PubMed Central identifications (PMCIDs) are available in Supplemental Table S3.

### External datasets

Human bulk liver organoid RNA-seq data (GSE215785 and GSE129457) and mouse bulk and single-cell liver developmental RNA-seq data (GSE90047) were downloaded from the Gene Expression Omnibus repository (44, 45, 47).

### Uploaded data and code availability

Nextflow code, BED files, and a Singularity container of software used for variant calling with ABRA2/Cadabra/Strelka2 are available on Zenodo (DOI: 10.5281/zenodo.10175692, https://zenodo.org/doi/10.5281/zenodo.10175692). R code for generating all figures using data from the supplemental tables is also available on both Zenodo and the GitHub project repository for this manuscript (https://github.com/isturgill/Sturgill_BAP1).

### Supplementary Data statement

Supplementary Data are available at NAR Online.

## Acknowledgements

We would like to thank David Marron and the UNC Lineberger Bioinformatics Core for assistance in implementing the somatic variant calling pipeline in Nextflow. We would also like to thank Peyton Kuhlers for expert feedback on statistics and modeling approaches. This research was supported by the National Cancer Institute award U24 CA264021 (Hoadley) and The National Institute of General Medical Sciences award R35GM147286 (Raab). IRS was supported by a UNC Bioinformatics and Computational Biology Training Fellowship.

## Author contributions

**IRS**: Data curation, formal analysis, investigation, software, visualization, writing – original draft. **JRR**: Funding acquisition, supervision, writing – original draft. **KAH**: Conceptualization, funding acquisition, supervision, writing – original draft. All authors read and approved the final manuscript.

## Supplemental Figures

Supplemental Figure S1. *BAP1* variant lengths for variants detected across mutation calling pipelines, comparing the prior TCGA MC3 dataset (n=182 variants) and the new dataset (n=257 variants) which combined updated TCGA GDC variant calls with calls from an ABRA2/Cadabra/Strelka2 workflow. Blue dotted line indicates the threshold for variant lengths ≥40bp.

Supplemental Figure S2. (A) Variant allele frequency for *BAP1* variants in pan-cancer mutant samples, separated by alteration type. P-value derived from two-sided Mann-Whitney U test with continuity correction. **(B)** Scatterplot of tumor purity and variant allele frequency for *BAP1* variants in pan-cancer mutant samples, colored by sample alteration type. Dotted line represents identity (y=x) line. Adjusted r-squared value derived from linear regression in R which performs QR decomposition followed by Wherry adjustment. Mut: mutation only, Mut+CN: mutation and gene-level copy number loss.

Supplemental Figure S3. Pan-cancer *BAP1* RNA-level expression by alteration type. P-values derived from pairwise two-sided Mann-Whitney U test with continuity correction and Bonferroni adjustment. CN: gene-level copy number loss only, Mut: mutation only, Mut+CN: mutation and gene-level copy number loss.

Supplemental Figure S4. Pan-cancer scatterplot of tumor *BAP1* RNA expression and *BAP1* activity score. Adjusted r-squared value derived from linear regression in R which performs QR decomposition followed by Wherry adjustment.

Supplemental Figure S5. Per-cancer boxplots of *BAP1* activity scores by alteration type. Un: unaltered, CN: gene-level copy number loss only, Mut+CN: mutation and gene-level copy number loss, Mut: mutation only. Red asterisk indicates statistical significance with p<0.05 from pairwise two-sided t-test with Bonferroni adjustment.

