## Supplemental Figures S1-S5 for "Expanded detection and impact of *BAP1* alterations in cancer"

### Supplemental Figure S1

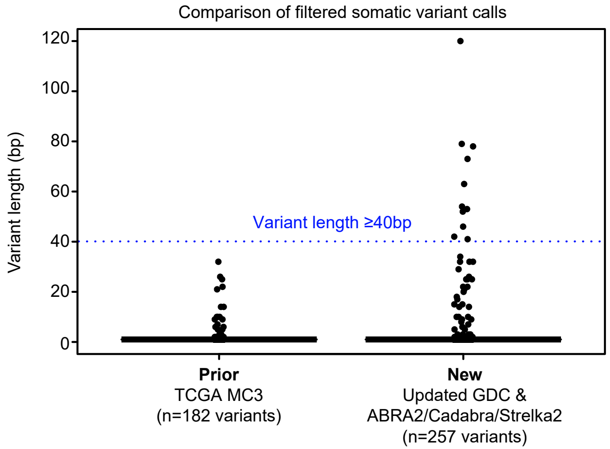

**Supplemental Figure S2**

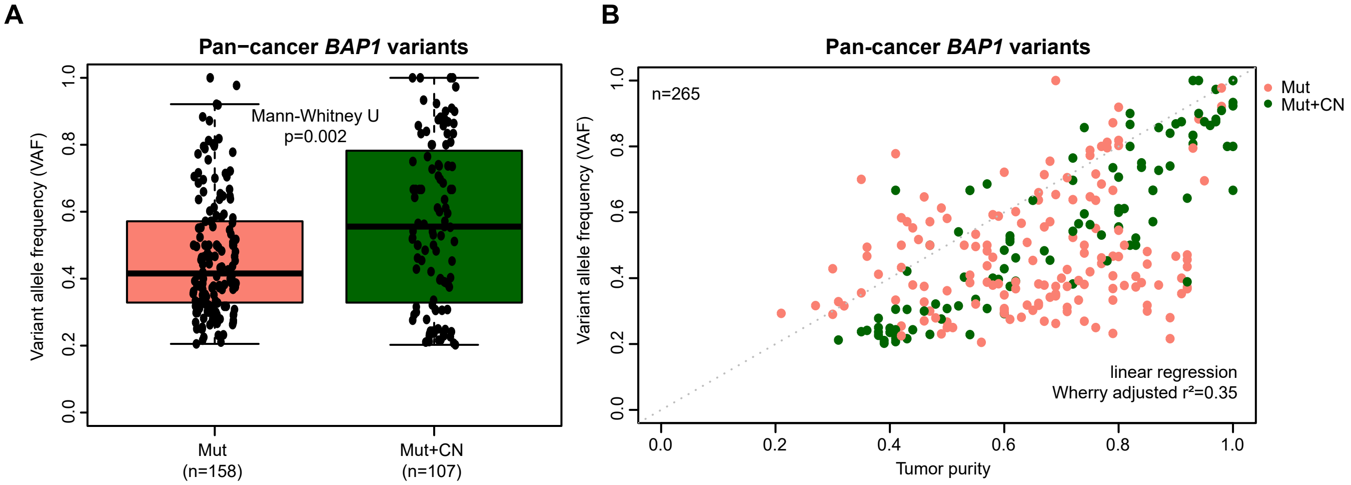

### Supplemental Figure S3

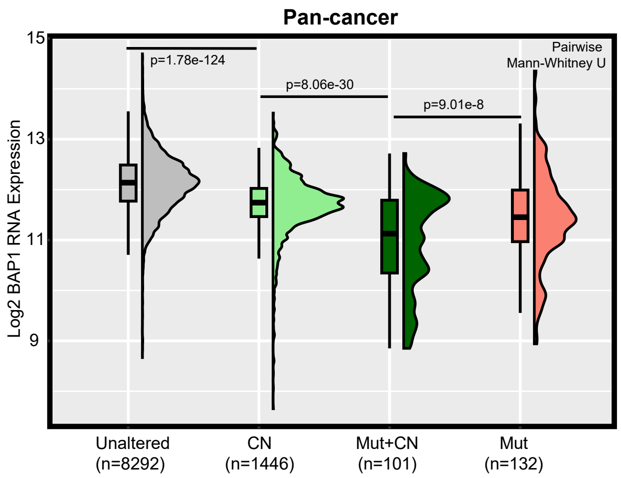

#### Supplemental Figure S4

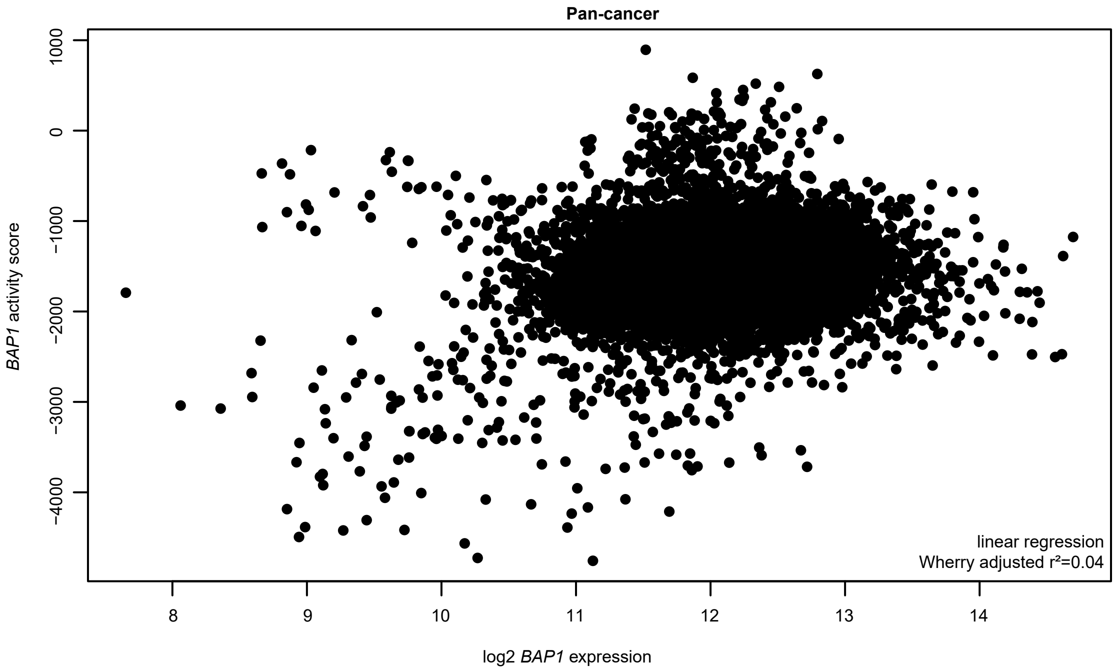

#### Supplemental Figure S5

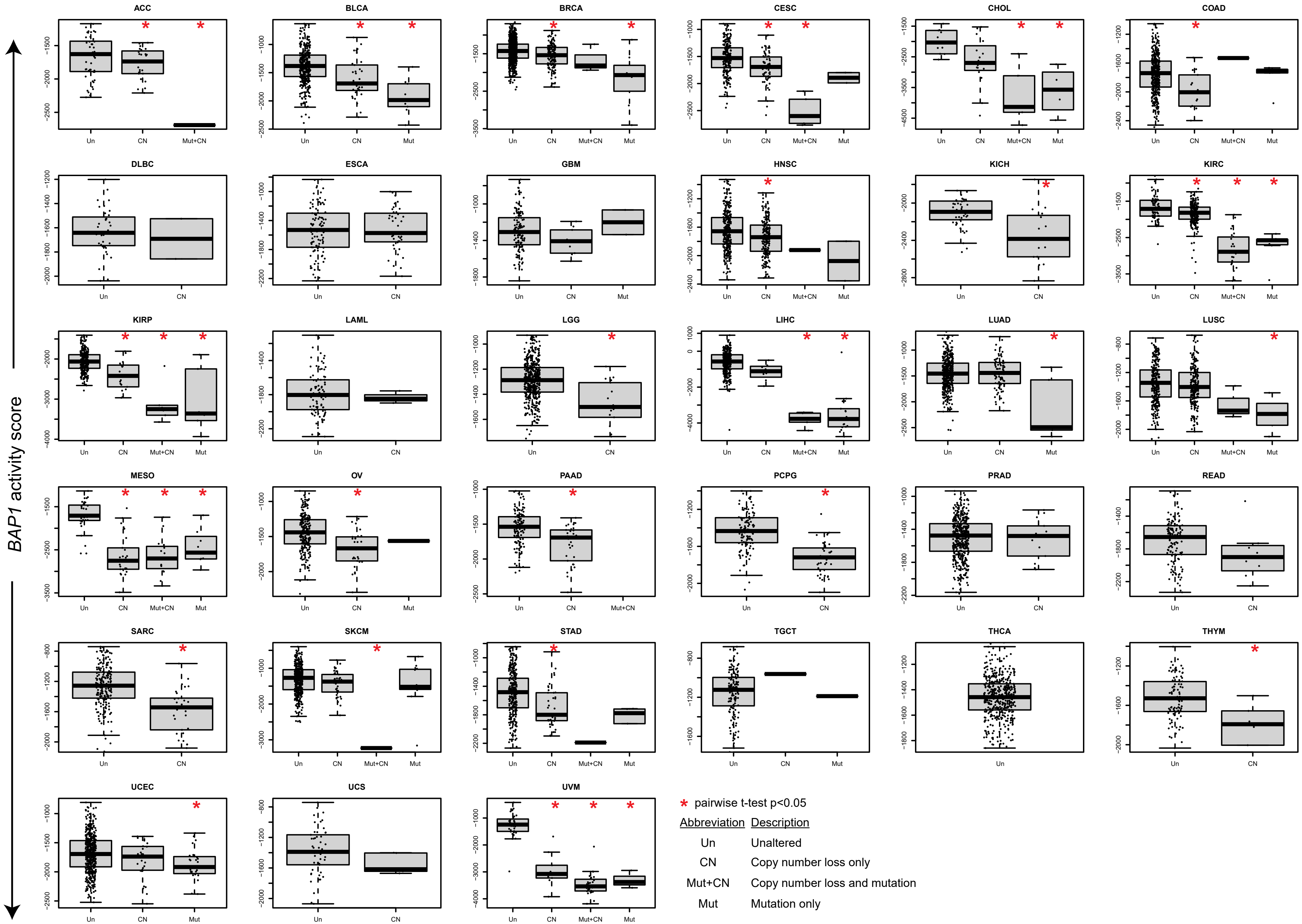
